# Identifying Priority Stepping Stone Reefs to Maintain Global Networks of Connected Coral Reefs

**DOI:** 10.1101/2025.08.30.673259

**Authors:** Ariel Greiner, Emily S. Darling, Amelia Wenger, Martin Krkošek, Marie-Josée Fortin

**Affiliations:** Department of Biology, University of Oxford, Oxford, UK; Department of Biology, Pennsylvania State University, University Park, PA, USA; Marine Program, Wildlife Conservation Society, Bronx, New York, USA; Department of Ecology and Evolutionary Biology, University of Toronto, Toronto, Ontario, Canada; Centre for Biodiversity and Conservation Science, School of the Environment, University of Queensland, St. Lucia, Queensland, Australia

**Keywords:** coral reef, conservation planning, marine, connectivity, metapopulations, stepping stone, network, biogeography

## Abstract

Conserving coral reef climate refugia is a key conservation strategy to address climate change. Yet, the accelerating impacts of coral bleaching and other anthropogenic pressures can jeopardize refugia persistence. The dispersal of coral larvae can increase long-term reef persistence through metapopulation dynamics if connected reefs can share beneficial adaptations to bleaching and facilitate demographic recovery. Here, we modelled reef connectivity using a network approach to identify present-day coral larval dispersal networks of predicted climate refugia and then determined locations of ‘stepping stone’ reefs that can be used to connect these global networks of refugia. We identify 10 key locations of coral reef stepping stones in Indonesia, Mozambique, Glorioso Islands, and Malaysia that may ensure that 84,564km^2^ of refugia could remain connected together even if other (non-refugia) reefs become degraded. Global coral reef conservation efforts should consider prioritizing these stepping stones to build more redundancy and resilience into future-proofing conservation strategies.

## Introduction

Coral reefs are a vital source of sustenance and livelihood for hundreds of millions of people worldwide (Cruz-Trinidad et al., 2014; FAO, 2012; Sing Wong et al., 2022), yet they face extreme threats from numerous anthropogenic pressures. Some of these pressures are global— such as climate change, which increases the frequency and severity of coral bleaching, cyclones and ocean acidification—while others vary locally, depending on the intensity of pressures like invasive species, overfishing, industrial development, and tourism (Andrello et al., 2022; Burke et al., 2011; Smith et al., 2016; Wenger et al., 2016). Although overfishing and water pollution are among the leading local pressures globally, the dominant pressures on reefs vary across regions (Andrello et al., 2022). By 2100, an estimated 70–99% of reefs could face long-term degradation due to coral bleaching (Frieler et al., 2013; van Hooidonk et al., 2016). Under worst-case climate scenarios, the long-term persistence of coral populations may depend on maintaining connectivity among a fragmented mosaic of climate-resilient habitat fragments (a metapopulation or network of reefs), supported by effective management of local pressures.

Predicting potential climate refugia for reefs has been considered in multiple regional studies (DeCarlo, 2020; McClanahan & Azali, 2021; McManus et al., 2020), but only Beyer et al. (2018) have considered a worldwide portfolio. Beyer et al. (2018) used historic, present-day and future projected thermal conditions, cyclone wave damage, as well as emigration of coral larvae simulation data to define 83 bioclimatic units (hereafter, ‘Refugia’) for coral reefs. Each of the Beyer et al. (2018) Refugium contain ∼500km^2^ of spatially contiguous coral reef cover that is predicted to be more resilient to thermal stress and cyclones and be good sources of coral larvae for recovery (due to their spatial position) (Beyer et al., 2018).

Since both global and local anthropogenic pressures are spatially variable, coral recovery can be achieved by coral larval dispersal among a network of reefs. Coral larvae dispersing from one reef to another connect coral reefs into distinct networks (‘reef networks’). Here, we define reef networks as the complete set of reef cells (18×18km geographic area containing coral cover, as defined based on the Global Distribution of Coral Reefs Dataset (UNEP-WCMC, WorldFish Centre, WRI, & TNC, 2010) and updates from the NOAA Coral Reef Watch Stations as derived from Wood et al. (2014), Beyer et al. (2018)) that send or receive coral larvae to/from each other according to a reference coral larval dispersal connectivity matrix (Greiner et al., 2022a). Using this empirically consistent reference coral larval dispersal connectivity matrix, Greiner et al. (2022a) found that many reef networks are quite large in size and span large regions (e.g. >50% of reef cells worldwide are in 6 reef networks). Thus, motivating coral reef conservation initiatives to consider these large spatial scales when determining which reefs to prioritize. In this study, we aim to identify reef cells that can act as ‘stepping stones’. Stepping stones are ecosystem patches (reef cells, in this case) that provide dispersal linkages between prioritized sets of ecosystem patches (in this case, the Refugia, which each have 2-157 reef cells), connecting or enhancing the connections between these prioritized ecosystem patches (Saura et al., 2014). Prioritizing stepping stones alongside prioritized ecosystem patches is essential for enhancing connectivity and achieving Target 3 of the Kunming-Montreal Global Biodiversity Framework (protecting 30% of terrestrial, inland water areas, coastal and marine areas), as well as for maintaining ecosystem function globally (Convention on Biological Diversity, 2022). While the stepping stone concept is general, stepping stones have mainly been determined for terrestrial ecosystems to date (Luo et al., 2021; Saura et al., 2017; Xu et al., 2024; but see D’Aloia et al., 2019). Identifying stepping stones for coral reefs will enable us to prioritize the protection or restoration of locations that could enhance connectivity between Refugia and provide spatial rescue through time (Gotelli, 1991; Mouquet & Loreau, 2003; Saura et al., 2014), and ideally provide crucial redundancy against the degradation of Refugia reef cells.

In this study, we determine which reef cells outside of Refugia (non-refugia reef cells) could act as stepping stones between Refugia to help ensure the long-term persistence of Refugia reef cells. We identify these ‘stepping stones’ through a network modelling approach based on a worldwide coral larval connectivity model (Greiner et al., 2022a). Specifically, we determine (1) a set of non-refugia reef cells (‘stepping stones’) that could, if prioritized for protection, maintain connections between Refugia reef cells when all other non-refugia reef cells are degraded; (2) the extent to which the stepping stones maintain connections among Refugia reef cells; and (3) the present-day anthropogenic pressure and protection status of the stepping stones.

## Methods

### Overview

We identify these stepping stones through a three step process: first, we determined the present-day reef networks of the 83 Refugia (Fig.1a); second, we simulated the degradation of all non-refugia reef cells by removing all non-refugia reef cells, and calculating the degraded reef networks of the 83 Refugia (Fig.1b); third, we used simulations to identify which non-refugia reef cells, if maintained, would restore connections between Refugia (i.e. act as ‘stepping stones’; Fig.1c). By doing so, we determined how many Refugia the stepping stones could help maintain connections with and the present-day status of those stepping stones. We simulate the degradation of all non-refugia reef cells to determine which non-refugia reef cells, if prioritized for protection now, could help maintain connections among the Refugia if only the Refugia remain. We envision that if the identified stepping stones are prioritized for protection at present, the coral reefs within them will have a better chance at withstanding future climatic or anthropogenic stressors than they would have had otherwise. We calculated all network designations and the present-day status of those stepping stones using R (v 4.3.1, R Core Team 2019) and QGIS (QGIS Development Team 2021).

**Figure 1.**
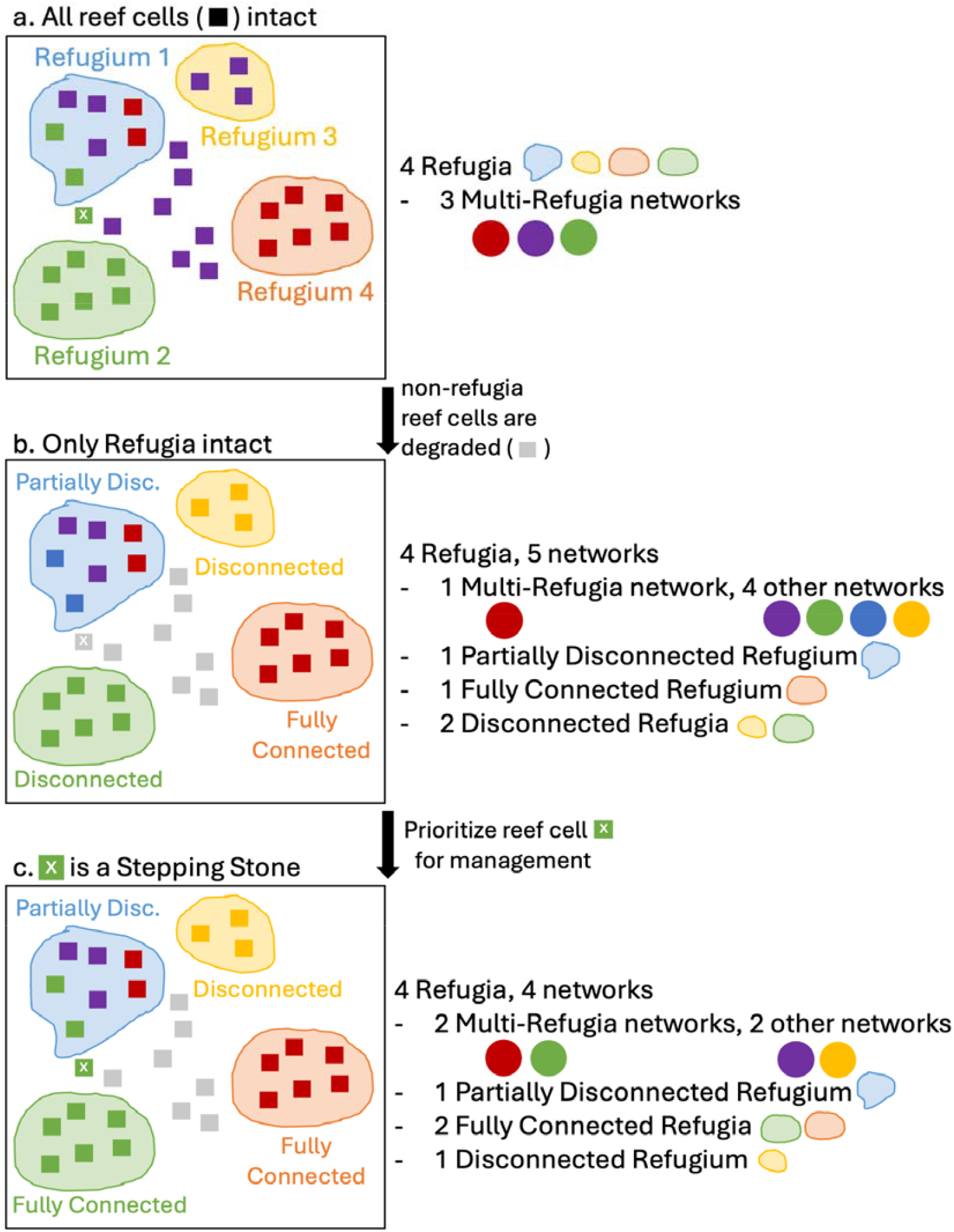
Method for Finding Stepping Stones. Reef cells (squares) in the same network are represented by the same coloured squares. Refugia are represented by large circles and degraded reefs are represented by grey squares (in (b) and (c)). When the ‘x’ non-refugia reef cell is added back in (c), through prioritizing reef cell ‘x’ for continued management, Refugia 1 and 2 are in the same reef network again. Note that this figure demonstrates that many reef cells may be included in one Refugium, that each Refugium may be part of multiple networks and that multiple Refugia may be in the same reef network.

### Reef Networks of the Refugia

To calculate the present-day reef networks of the 83 Refugia, we used the connectivity matrix generated by Greiner et al. (2022a) that was derived from Wood et al. (2014). First, we used the Andrello et al. (2022) grid cells (which used data from Beyer et al. (2018)) to determine which of the Wood et al. (2014) grid cells (hereafter, ‘reef cells’) were in Refugia (hereafter, ‘Refugia reef cells’). We work at the level of reef cells because that is the spatial resolution for which we have connectivity information for reefs worldwide from Greiner et al. (2022a). In this study, we will use the ‘reference connectivity matrix’ generated in Greiner et al. (2022a) (see SM1 for more details).

First, we calculated which Refugia reef cells are in which reef networks (‘weak cluster’ in igraph in R; Csárdi et al. 2023) and then determined which Refugia reef cells were in the same reef networks as other Refugia reef cells and which were not in the same reef networks as any other Refugia reef cells (SM2-Fig.1). Note that based on how the Refugia reef cells were identified by Beyer et al. (2018), they should export more larvae to other reef cells than the average reef cell. However, this does not necessarily imply that a large proportion of those receiving reef cells are other Refugia reef cells.

Then, we removed all of the non-refugia reef cells and recalculated the reef networks of the 83 Refugia by removing the rows and columns of the connectivity matrix that corresponded with the non-refugia reef cells. By doing this, we determined which Refugia would remain connected to other Refugia (‘Fully Connected Refugia’), which Refugia would lose some connections with other Refugia (‘Partially Disconnected Refugia’), and which would lose all connections with other Refugia (‘Disconnected Refugia’) if all the non-refugia reef cells were degraded (see SM3-Fig.1).

### Identifying and Characterizing Stepping Stones

To identify stepping stones, we focused on the Partially Disconnected Refugia and the Disconnected Refugia (e.g. Refugia 1, 2, and 3 in Fig.1b; reefs in SM3-Fig.1b,c). First, we examined non-refugia reef cells that were part of the same networks as the Disconnected Refugia before any reef cells were removed. We then checked whether these non-refugia reef cells had direct connections—either sending or receiving larvae, or both—with at least one reef cell in a Disconnected Refugium (i.e. ‘Disconnected Refugium *i*’). Finally, we determined whether these non-refugia reef cells were also directly connected to a reef cell from any other Refugia that were originally in the same network as Disconnected Refugium *i*. Any reef cell that satisfied both of these conditions was defined as a stepping stone. We distinguished which of those stepping stones were ‘connector’ stepping stones or ‘sink’ stepping stones, depending on whether the stepping stones received larvae from one Refugium and sent larvae to at least one other Refugium (connector) or received larvae from multiple Refugia but did not send larvae to any Refugia (sink). We performed this analysis for every Disconnected Refugium. We then repeated the process for non-refugia reef cells connected to Partially Disconnected Refugia to find additional stepping stones.

Once the connector and sink stepping stones had been identified, we evaluated how adding them back affected the connectivity of the Refugia reef networks. Specifically, we counted how many Refugia become (1) fully re-connected with the Refugia they were connected to in the present-day networks (i.e., become ‘Fully Connected Refugia’), (2) remain or become Partially Disconnected Refugia, and (3) remain Disconnected Refugia (SM3-Fig.2). To better indicate the scale of these changes, we also noted the total number of Refugia reef cells and the total reef cell area that changed category. Note that ‘total reef cell area’ corresponds to the total size of the reef cells themselves and is an overestimate of the area within those cells that contains coral cover itself. We could not calculate total coral cover itself because precise estimates of the percent coral cover was not available for all of the Refugia reef cells. Lastly, we used data from Andrello et al. (2022) to assess which local anthropogenic pressures were affecting each stepping stone and the protected planet database (UNEP-WCMC and IUCN, 2024) to evaluate the protection status of each stepping stone. From the analysis of Andrello et al. (2022), we gathered the top anthropogenic pressure level and the cumulative anthropogenic pressure level of each stepping stone (see Fig.4).

**Figure 2.**
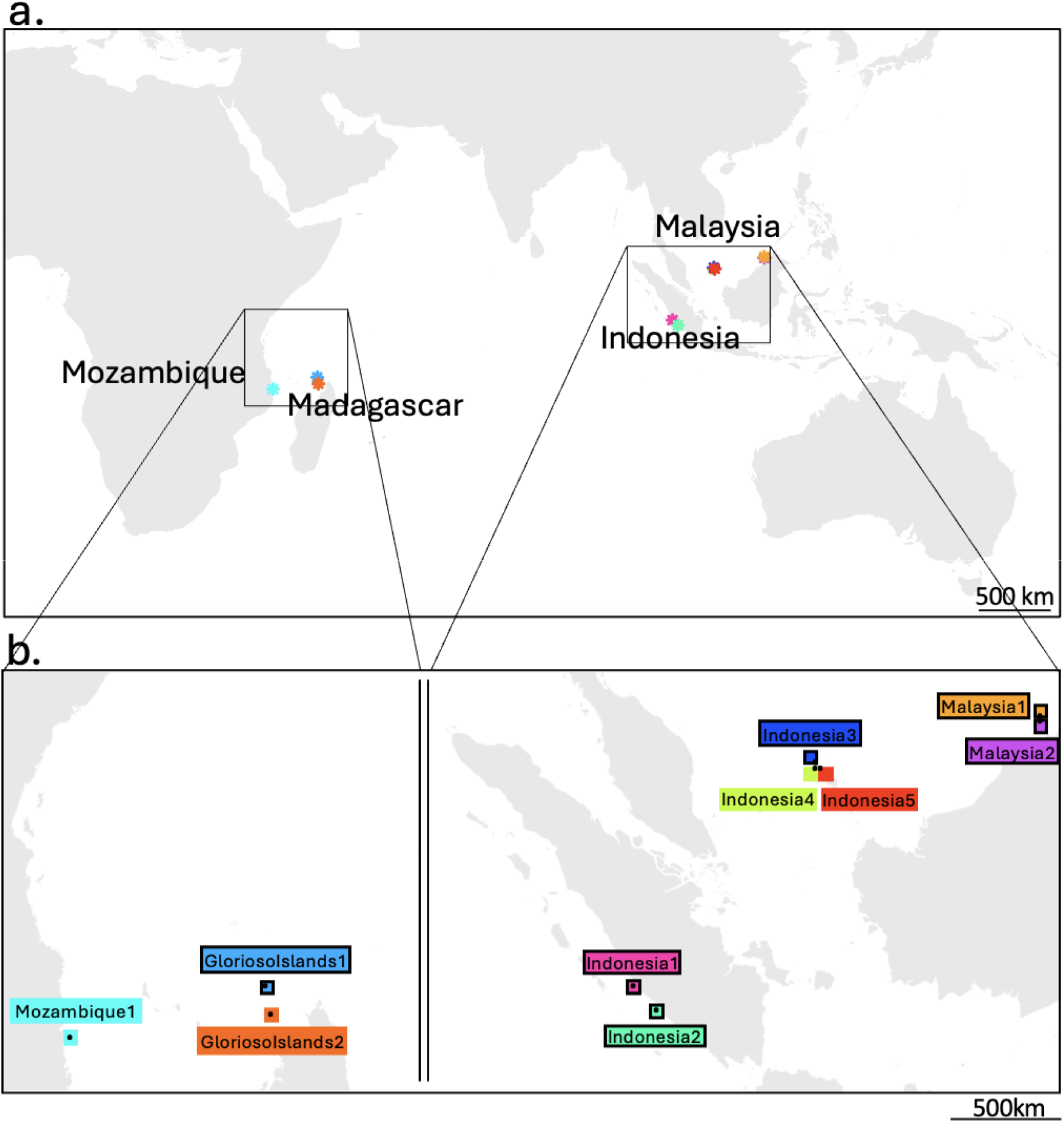
Stepping Stones. (a) Each point represents a different stepping stone reef cell. (b) Black dots represent the spatial extent and precise locations of stepping stone reef cells. The coloured borders around each stepping stone distinguish them from each other. Black borders in (b) indicate the ‘connector’ stepping stones, while the remainder are the ‘sink’ stepping stones. Land polygons in (b) are from naturalearthdata.com (10m scale resolution, accessed August 19, 2025).

## Results

### Stepping Stones

We identified ten non-refugia reef cells as stepping stones (Fig.2). These ten stepping stones were the only non-refugia reef cells that could maintain connections between Refugia following a scenario when all other non-refugia reef cells were degraded. Six of these stepping stones are ‘connector’ stepping stones and four of them are ‘sink’ stepping stones; these ten stepping stones are located in Indonesia, Mozambique, Glorioso Islands, and Malaysia (Fig.2).

### Refugia Connectivity with and without Stepping Stones

When all non-refugia reef cells are degraded, the reef networks become more fragmented and less connected (Fig.3a,b; SM3-Fig.1). In particular, the number of Refugia in multi-Refugia networks (networks that contain Refugia reef cells from more than one Refugia) decreases from 73 (3,760 reef cells, 1,218,240km^2^ total reef cell area) to 66 (3,388 reef cells, 1,097,712km^2^) and the average reef network size declines from 25,364.57km^2^ to 11,548.1km^2^. (median network size is unchanged at 648km^2^). Nineteen Refugia maintain all connections with other Refugia, 47 Refugia lose some of their connections with other Refugia (Partially Disconnected Refugia), and seven Refugia lose all connections with other Refugia (Disconnected Refugia) (Fig.3b; SM3-Fig.1).

**Figure 3.**
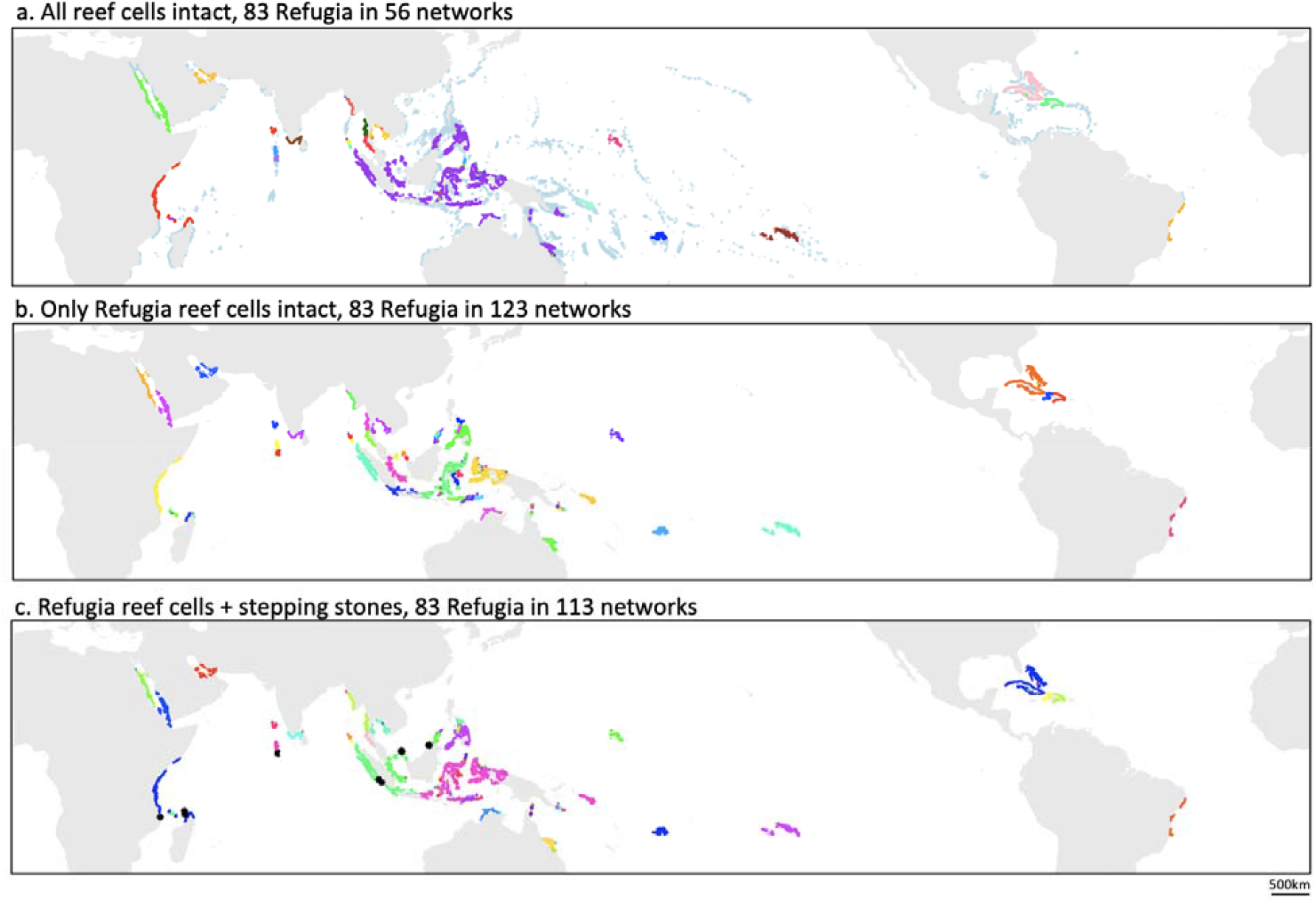
Re-Connecting 83 Refugia Networks Worldwide. Bright coloured (i.e. non-light blue) reef cells indicate Refugia reef cells, coloured according to which reef network they are in (same reef network, same colour). (a) The light blue reefs indicate reef cells that are not in Refugia. (c) Black stars indicate stepping-stone reef cells. Note that if only the ‘connector’ stepping stones are added back in, the 84 Refugia are in 114 networks (Supplementary Materials 3 - Fig. 3).

When the 10 stepping stone reef cells are added back in (3240km^2^, Fig.2), the average network size increases to 12,598.73km^2^ (median remains unchanged at 648km^2^) and 69 Refugia are now in multi-Refugia networks (3,501 reef cells, 1,134,324km^2^) (Fig.3c; SM3-Fig.2). Specifically, nine Refugia (294 grid cells, 95,256km^2^) become more connected to other Refugia – 8 Refugia (261 grid cells, 84,564km^2^) become Fully Connected Refugia (6 Partially Disconnected to Fully Connected Refugia, 2 Disconnected to Fully Connected Refugia) and 3 Refugia (113 grid cells, 36,612km^2^) become connected to at least one other Refugia again (2 Disconnected Refugia to Fully Connected Refugia and 1 Disconnected Refugium to a Partially Disconnected Refugium) (SM3-Fig.2). If only consider the 6 ‘connector’ stepping stone reef cells (1944km^2^, Fig.2), the average network size becomes 12,476.84km^2^ (median remains unchanged at 648km^2^) and 3 Refugia (113 grid cells, 36,612km^2^) become connected to at least one other Refugium again (Disconnected to Partially Disconnected Refugia), so the number of Refugia in multi-Refugia networks remains the same (69 Refugia; 3,501 reef cells, 1,134,324km^2^). However, with only the 6 ‘connector’ stepping stones, no additional Refugia become Fully Connected Refugia (i.e. none of the Partially Disconnected Refugia become Fully Connected Refugia, none of the Disconnected Refugia become Partially Disconnected Refugia) (SM3-Fig.4).

**Figure 4.**
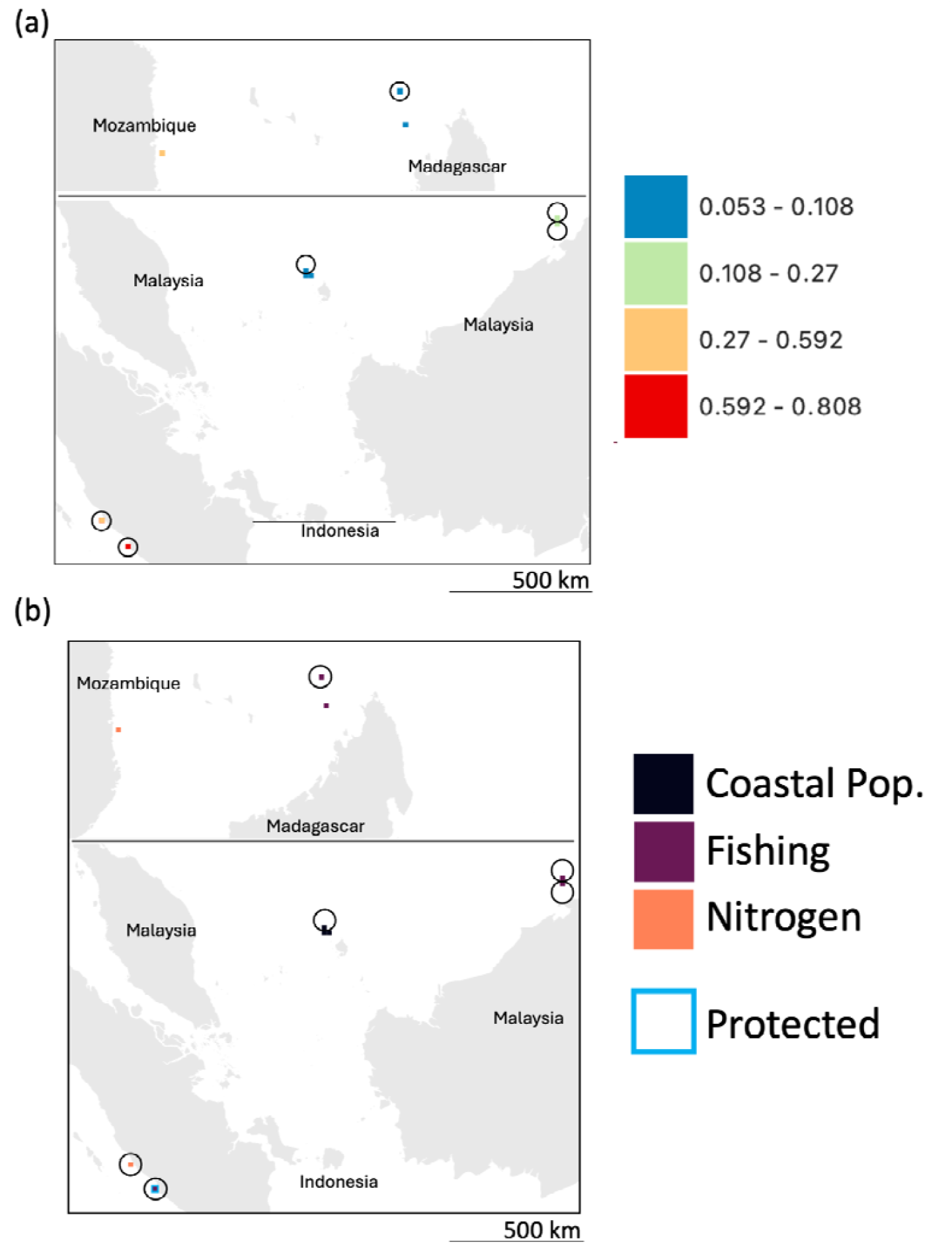
Anthropogenic Pressure Scores of Stepping Stones. The colour of each stepping stone indicates their (a) cumulative anthropogenic pressure score or their (b) top anthropogenic pressure as calculated by Andrello et al. (2022). To calculate these metrics, Andrello et al. (2022) calculated the percentile of each pressure (Fishing, Coastal Population, Industrial Development, Tourism, Sediments, Nitrogen) within a reef cell from the global distribution of each pressure. The top anthropogenic pressure level of each stepping stone is the pressure with the highest percentile for that stepping stone. The cumulative anthropogenic pressure level is the average of the percentiles of the six pressure layers within that reef cell (additional details in Supplementary Materials 1). Black circles around the stepping stones indicate the ‘connector’ stepping stones, while the remainder are ‘sink’ stepping stones. Land polygons are from naturalearthdata.com (10m scale resolution, accessed August 19, 2025). (a) The colour of the stepping stone reef cells indicates their cumulative anthropogenic pressure score. Cumulative anthropogenic pressure scores were derived from Andrello et al. (2022)’s scores and range between 0 and 1. (b) The colour of each stepping stone reef cell indicates the top anthropogenic pressure according to Andrello et al. (2022) experienced by each stepping stone. Five stepping stones are most threatened by high fishing levels, two by high nitrogen levels and three by proximity to large coastal populations. The blue border indicates the stepping stone that is protected according to the World Database on Protected Areas (accessed on April 3, 2024). Note that because of imperfect overlap between the grid cells of Andrello et al. (2022) and this study (those from Wood et al. 2014), some of the stepping stones overlapped with multiple of Andrello et al. (2022)’s grid cells–though this only caused a discrepancy with one of them (blue-bordered stepping stone; 5 fishing grid cells, 1 industrial development grid cell and 1 coastal population grid cell, see Supplementary Materials 4).

### Properties of the Stepping Stones

The stepping stones varied in cumulative anthropogenic pressure level (0-1, 1 indicating high pressure), with the stepping stones near South Sumatra experiencing the highest cumulative pressure levels and the stepping stones near Northern Madagascar and the Riau Islands experiencing the lowest cumulative pressure levels (Fig.4a). Only one stepping stone is within a protected or managed area identified by the World Database on Protected Areas (UNEP-WCMC and IUCN, 2024; Pantai Panjang dan P.Baai nature recreation park) (Fig.4b). The stepping stones vary in the top pressure that they experience (Andrello et al. 2022; Fig.4b).

## Discussion

By prioritizing the reefs within the 10 stepping stones for conservation management, we maintain Refugia-Refugia connections with nine Refugia that would otherwise be disconnected with the degradation of non-refugia reef cells. In doing so, these 10 stepping stones may indirectly enhance the resilience of reefs covering an area up to 25 times larger than the prioritized region itself (3240km^2^ vs 84,564km^2^, Fig.3c). These stepping stones are especially important because they are the only individual non-refugia reef cells capable of independently maintaining connectivity between Refugia. This small number reflects the fact that many Refugia remain within multi-Refugia networks even when only Refugia reef cells are intact (Fig.3b; as suggested in Greiner et al. 2022a), indicating that the Refugia alone may already provide some level of spatial rescue and redundancy to themselves. Note that Beyer et al. (2018) accounted for connectivity in their scoring by prioritizing reef cells that exported many larvae to other reef cells. Thus, it is unsurprising that the reef cells within the Refugia are well-connected to other reef cells, but it was not clear *a priori* that they would be well-connected to reef cells in the same or other Refugia specifically. Six of these 10 are an even higher priority for conservation as they are connector stepping stones, and may allow larvae to flow from one Refugia to another (Fig.2). By only prioritizing these 6, the same number (69) of Refugia are restored to multi-Refugia networks but none of the Refugia become fully re-connected to other Refugia (SM3-Fig.3,4). It is hopeful that seven of the 10 stepping stones have a cumulative anthropogenic pressure score below 0.5 (Fig.4a), indicating that they may have relatively few non-climate pressures. However, monitoring is needed to confirm the ecological integrity or capacity of these locations to have functioning coral communities that can promote coral larval dispersal. Only one of the stepping stone reef cells is currently designated as ‘protected’ (Fig.5); notably, this is also the cell with the highest cumulative anthropogenic pressure score (Fig.4), highlighting a fortunate alignment between protection status and conservation urgency (even if protection is unlikely to protect against all pressures).

These 10 candidate stepping stones were chosen based on present-day knowledge of the location of reefs, ocean currents and coral larval dispersal. The goal of this study was to use what we know of global coral larval dispersal connectivity to identify possible reefs that, if individually prioritized for protection, could help maintain the 83 Refugia (Beyer et al. 2018). We searched for single reef cell stepping stones because a single reef cell is already quite a large region to prioritize and we wanted to find the least additional reef area to prioritize (note that pairs of reef cells did not re-connect any Refugia, see SM5). We focus on these 83 Refugia as they currently guide how various reef conservation initiatives and funders (e.g., the Global Fund for Coral Reefs, Bloomberg Ocean Initiative) prioritize reefs for protection around the world, as they are thought to have the best chance of being resilient towards future climate impacts. That being said, this study is meant as a starting point for this type of work and more specific analyses should be performed in the regions containing the 10 stepping stones identified here to determine which reefs within these reef cells are most relevant and practical to prioritize (as there are unlikely to be 3,240km^2^ of reefs within these stepping stones). In particular, it would be helpful to collect species-specific coral larval dispersal information and perform genetic analyses on coral from the reefs found in these 10 stepping stones (and for reefs in general) to ascertain whether these reefs do send and/or receive larvae from reefs in reef cells of more than one Refugia–as the data that we are using to find these stepping stones is consistent with empirical data (see SM1 and Wood et al. 2014 and Greiner et al. 2022a for more information), has been used to drive coral reef conservation priorities (see Beyer et al. 2018) and is the only global connectivity matrix for coral that is available (at this time), but is not directly derived from empirical data.

This study presents a methodology for identifying candidate stepping stone reef cells that can be adapted for other at-risk ecosystems and used to guide prioritization decisions when connectivity is a management goal. For example, this methodology could be used to identify key stepping-stone reefs at regional scales. It could also be used for other at-risk marine ecosystems whose key organisms disperse passively along ocean currents (e.g. rocky reefs). One could also calculate candidate stepping stone reef cells that focus on reef fish larval dispersal networks; it would be interesting to determine whether similar reef cells would be prioritized and how well connected the Refugia are from a reef fish perspective. Lastly, it would be interesting to assess whether any of these stepping stones may also be likely to send macroalgal gametes between Refugia, as macroalgal gamete dispersal may counteract the coral larval dispersal and tip reefs in these Refugia to a degraded, less coral-dominated state (see Greiner et al., 2022b). The risk of reefs tipping into a degraded, less coral-dominated state may become more relevant as coral reefs become more disconnected, as dispersal between reefs has been shown to stabilize reefs (see Greiner et al. 2025). In general macroalgal gametes are not thought to disperse as far as coral larvae (Deysher and Norton, 1981; Mumby, 2006; Stiger and Payri, 1999), but long distance dispersal of macroalgal gametes along the same ocean currents that transport coral larvae is possible and ensuring that the reefs in those reef cells maintain low macroalgal cover might be crucial for preserving high coral cover in connected reefs. Comparing which reef cells are identified as crucial stepping stones for all of these different taxa could further help target coral reef prioritization efforts around the world and allow us to see which reefs may be most efficient to prioritize to satisfy multiple different conservation priorities.

## Conclusion

In this study, we identify 10 stepping stones capable of maintaining connections among Refugia reef cells. The reefs within these stepping stones are exposed to a range of anthropogenic pressures, and only one currently holds protected status. The reefs within these 10 stepping stones should be prioritized for conservation and local management before further degradation occurs, as our study identifies these locations as the only individual reef cells that can maintain connectivity between Refugia at this spatial scale (according to the analysis in this study). Preserving healthy reefs in these stepping stones will enhance the resilience of reefs in nine Refugia. Strengthening the resilience of Refugia reef cells increases the likelihood that these reefs—predicted to have the greatest potential to maintain coral cover under climate change— will persist and continue supplying coral larvae to other reefs, thereby supporting protection of a much larger global reef area. These findings also underscore the need for further research into larval and gamete connectivity across reef systems to better identify key connectors (across spatial scales) and thus optimize global coral reef conservation strategies.

## Acknowledgements

We thank Marco Andrello for his helpful comments and Cole B. Brookson for a helpful R function for map generation.

## Data Availability Statement

The climate refugia scores and pressure scores for each Andrello et al. (2022) reef cell can be found at https://github.com/WCS-Marine/local-reef-pressures. The code used to generate the connectivity matrix used in this study can be found in https://github.com/ArielGreiner/GlobalCoralConnectivity_SpatialRescue_Project, while the original world-wide coral reef connectivity matrix generated for Wood et al. (2014) and the grid cell coordinates are available at the University of Bristol data repository, data.bris, at: https://doi.org/10.5523/bris.2s0fn0bc89omq2kj2rol7iolwt (Hendy & Wood, 2022). The scripts used to modify these datasets and perform the analyses described above and the datasets needed for doing so are available at: https://figshare.com/s/3d6dc00129a1da24820c.

## References

Andrello, M., Darling, E. S., Wenger, A., Suárez□Castro, A. F., Gelfand, S., & Ahmadia, G. N. (2022). A global map of human pressures on tropical coral reefs. Conservation Letters, 15(1), e12858.

Beyer, H. L., Kennedy, E. V., Beger, M., Chen, C. A., Cinner, J. E., Darling, E. S., … & Hoegh□Guldberg, O. (2018). Risk□sensitive planning for conserving coral reefs under rapid climate change. Conservation Letters, 11(6), e12587.

Burke, L., Reytar, K., Spalding, M., & Perry, A. (2011). Reefs at risk revisited: technical notes on modeling threats to the world’s coral reefs. Washington, DC: World Resources Institute.

Convention on Biological Diversity. (2022). Decision 15/4: Kunming-Montreal Global Biodiversity Framework. CBD/COP/DEC/15/4

Cruz-Trinidad, A., P. M. Aliño, R. C. Geronimo, and R. B. Cabral. 2014. “Linking food security with coral reefs and fisheries in the coral triangle.” Coastal Management 42:160–182.

Csárdi G., T. Nepusz, V. Traag, S. Horvát, F. Zanini, D. Noom, and K. Müller. 2023. “Igraph: Network Analysis and Visualization in R.” doi:10.5281/zenodo.7682609

D’Aloia, C. C., Naujokaitis-Lewis, I., Blackford, C., Chu, C., Curtis, J. M., Darling, E., … & Fortin, M. J. (2019). Coupled networks of permanent protected areas and dynamic conservation areas for biodiversity conservation under climate change. Frontiers in Ecology and Evolution, 7, 27.

DeCarlo, T. M. (2020). Treating coral bleaching as weather: a framework to validate and optimize prediction skill. PeerJ, 8, e9449.

Deysher L, Norton TA (1981) Dispersal and colonization in Sargassum muticum (Yendo) Fensholt. J Exp Mar Biol Ecol 56:179–195. 10.1016/0022-0981(81)90188-X

FAO. 2012. “The State of World Fisheries and Aquaculture.” FAO Fisheries and Aquaculture Department, Food and Agriculture Organization of the United Nations, Rome. - SOFIA 2012

Frieler, K., Meinshausen, M., Golly, A., Mengel, M., Lebek, K., Donner, S. D., & Hoegh-Guldberg, O. (2013). Limiting global warming to 2°C is unlikely to save most coral reefs. Nature Climate Change, 3(2), 165–170.

Gotelli, N. J. (1991). Metapopulation models: The rescue effect, the propagule rain, and the core-satellite hypothesis. The American Naturalist, 138(3), 768–776.

Greiner, A., Andrello, M., Darling, E., Krkošek, M., & Fortin, M. J. (2022a). Limited spatial rescue potential for coral reefs lost to future climate warming. Global Ecology and Biogeography, 31(11), 2245–2258.

Greiner, A. S., Darling, E., Fortin, M. J., & Krkošek, M. (2022b). The combined effects of dispersal and herbivores on stable states in coral reefs. Theoretical Ecology, 15(4), 321–335.

Greiner, A., Andrello, M., Krkošek, M., Fortin, M. J., Nand, Y., Jupiter, S. D., … & Darling, E. S. (2025). Dispersal can spread management benefits: Insights from a modeled Fijian coral reef network. Ecological Applications, 35(8), e70156.

Luo, Y., Wu, J., Wang, X., & Peng, J. (2021). Using stepping-stone theory to evaluate the maintenance of landscape connectivity under China’s ecological control line policy. Journal of Cleaner Production, 296, 126356.

Manel, S., Loiseau, N., Andrello, M., Fietz, K., Goñi, R., Forcada, A., … & Mouillot, D. (2019). Long-distance benefits of marine reserves: myth or reality?. Trends in Ecology & Evolution, 34(4), 342–354.

McClanahan, T. R., & Azali, M. K. (2021). Environmental variability and threshold model’s predictions for coral reefs. Frontiers in Marine Science, 8, 778121.

McManus, L. C., Vasconcelos, V. V., Levin, S. A., Thompson, D. M., Kleypas, J. A., Castruccio, F. S., … & Watson, J. R. (2020). Extreme temperature events will drive coral decline in the Coral Triangle. Global Change Biology, 26(4), 2120–2133.

Mouquet, N., & Loreau, M. (2003). Community patterns in source-sink metacommunities. The American Naturalist, 162(5), 544–557.

Mumby PJ (2006) The impact of exploiting grazers (Scaridae) on the dynamics of Caribbean coral reefs. Ecol Appl 16:747–769. 10.1890/1051-0761(2006)016[0747:TIOEGS]2.0.CO;2

QGIS Development Team (2021). QGIS Geographic Information System. Open Source Geospatial Foundation Project.

R Core Team. 2019. “R: A language and environment for statistical computing.” R 1145 Foundation for Statistical Computing, Vienna, Austria. https://www.R-project.org/.

Saura, S., Bodin, Ö., & Fortin, M. J. (2014). EDITOR’S CHOICE: Stepping stones are crucial for species’ long□distance dispersal and range expansion through habitat networks. Journal of Applied Ecology, 51(1), 171–182.

Saura, S., Bastin, L., Battistella, L., Mandrici, A., & Dubois, G. (2017). Protected areas in the world’s ecoregions: How well connected are they?. Ecological Indicators, 76, 144–158.

Sing Wong, A., S. Vrontos, and M. L. Taylor. 2022. “An assessment of people living by coral reefs over space and time.” Global Change Biology 28:7139–7153.

Smith, J. E., Brainard, R., Carter, A., Grillo, S., Edwards, C., Harris, J., … & Sandin, S. (2016). Re-evaluating the health of coral reef communities: baselines and evidence for human impacts across the central Pacific. Proceedings of the Royal Society B: Biological Sciences, 283(1822), 20151985.

Stiger V, Payri CE (1999) Spatial and temporal patterns of settlement of the brown macroalgae Turbinaria ornata and Sargassum mangarevense in a coral reef on Tahiti. Mar Ecol Prog Ser 191:91–100. 10.3354/meps191091

UNEP-WCMC and IUCN (2024), Protected Planet: The World Database on Protected Areas (WDPA) and World Database on Other Effective Area-based Conservation Measures (WD-OECM) [Online], April 2024, Cambridge, UK: UNEP-WCMC and IUCN. Available at: https://www.protectedplanet.net.

UNEP-WCMC, WorldFish Centre, WRI, TNC (2010) Global distribution of coral reefs, compiled from multiple sources including the Millennium Coral Reef Mapping Project. Version 1.3. Includes contributions from IMaRS-USF and IRD (2005), MaRSUS (2005) and Spalding et al. (2001). Cambridge (UK): UNEP World Conservation Monitoring Centre. URL: http://data.unep-wcmc.org/datasets/1

Van Hooidonk, R., Maynard, J., Tamelander, J., Gove, J., Ahmadia, G., Raymundo, L., Williams, G., Heron, S. F., & Planes, S. (2016). Local-scale projections of coral reef futures and implications of the Paris Agreement. Scientific Reports, 6(1), 1–8.

Wenger, A. S., Williamson, D. H., da Silva, E. T., Ceccarelli, D. M., Browne, N. K., Petus, C., & Devlin, M. J. (2016). Effects of reduced water quality on coral reefs in and out of no□take marine reserves. Conservation Biology, 30(1), 142–153.

Wood, S., Paris, C. B., Ridgwell, A., & Hendy, E. J. (2014). Modelling dispersal and connectivity of broadcast spawning corals at the global scale. Global Ecology and Biogeography, 23(1), 1–11.

Xu, D., Peng, J., Jiang, H., Dong, J., Liu, M., Chen, Y., … & Meersmans, J. (2024). Incorporating barriers restoration and stepping stones establishment to enhance the connectivity of watershed ecological security patterns. Applied Geography, 170, 103347.

