## SupplementaryMaterials for "Identifying Priority Stepping Stone Reefs to Maintain Global Networks of Connected Coral Reefs"

**Supplementary Materials 1**. Extra Methods Details

*Determining whether a Reef Cell was in a ‘Refugia’*

The grid cells used by Beyer et al. (2018) were smaller than the Wood et al. (2014) grid cells (324km^2^) for which we had connectivity information and were not perfectly aligned so we had to use the spatial join function (in the sf package in R; Pebesma & Bivand, 2018; Pebesma 2018) to match the two sets of grid cells; if even one of the Andrello et al. (2022) Refugia grid cells was in a Wood et al. (2014) grid cell, that grid cell was labeled as a Refugia grid cell.

*Greiner et al. (2022a)’s Reference Connectivity Matrix*

Greiner et al., (2022a)’s reference connectivity matrix is based on the global connectivity matrix for a generic broadcast-spawning scleractinian coral larva generated in Wood et al. (2014), modified so that the median Euclidean dispersal distance was 42km (consistent with empirical findings in Manel et al. 2019) to make it more likely to contain coral larval dispersal links relevant for spatial rescue (Gotelli, 1991; Mouquet & Loreau, 2003; Saura et al., 2014) (see SM1). Note that, Greiner et al. (2022a)’s reference connectivity matrix found reef networks similar to those delineated in empirical studies (Cowen et al. 2006; Holstein et al. 2014) and preserved many of the similarities to empirical results of the connectivity matrix of Wood et al. (2014) (e.g. Ayre & Hughes, 2000; Kool et al. 2011; for more details see Greiner et al. 2022a).

*Additional Connectivity Assumptions*

We assume that all reef cells included in the Greiner et al., (2022a) reference connectivity matrix still contain at least some ‘not-degraded’ coral cover. Note that we consider two reef cells to be connected if they exchanged at least one larva—either sent or received—over the period from November 2003 to November 2011. This is based on simulated global coral larval dispersal data from Wood et al. (2014), as modified in Greiner et al. (2022a). This method assumes that, in general, coral larval connectivity in the future driven by ocean currents will not change at the spatial scale that we are considering; i.e. no new connections will form between reef cells that are not currently connected, and that existing connections will only be lost through loss of coral reef cover in reef cells (this is consistent with Vogt-Vincent et al., 2023). Our overall approach to finding stepping stones is based on the assumption that some of the connections between Refugia will be maintained through the continued maintenance of the coral reefs in the Refugia reef cells (through their present-day prioritization), while other connections between Refugia rely on non-refugia reef cells.

*Conservation Status and Exposure to Local Pressures of Potential Stepping Stones*

Once we had identified stepping stones, we assessed the conservation status of each stepping stone. We used data from Andrello et al., (2022) to determine how anthropogenic pressures were affecting each stepping stone, and the protected planet database (UNEP-WCMC and IUCN, 2024) to evaluate the protection status of the stepping stones. The Andrello et al. (2022) database contained information on which anthropogenic pressures affect grid cells containing coral reefs around the world, which anthropogenic pressure is the top pressure of said grid cells and then ranks all the grid cells in terms of their cumulative anthropogenic pressure level (estimated by combining pressure scores of each of the different pressures, see Andrello et al. (2022) for more details). The Andrello et al. (2022) database explores 6 anthropogenic pressures: fishing, coastal population, industrial development, tourism, sediments and nitrogen. We again used the spatial join function (in the sf package (Pebesma & Bivand, 2018; Pebesma 2018) in conjunction with QGIS) to determine which Andrello et al. (2022) grid cells overlapped with the stepping stone reef cells to determine the top anthropogenic pressure and cumulative anthropogenic pressure score of the Andrello et al. (2022) grid cells that overlapped with the stepping stone reef cells. We followed the same spatial join procedure to determine which EEZ each stepping stone was in, as the Andrello et al. (2022) database contained this information as well. To determine the legal protection status of each of the stepping stone reef cells, we downloaded the protected area shapefiles of Africa and Asia from protectedplanet.net (UNEP-WCMC and IUCN, 2024, accessed on April 3, 2024) and then used QGIS to compare overlap between known protected areas and the stepping stone reef cells.

All R code used to generate these results and these figures can be found in <https://figshare.com/s/3d6dc00129a1da24820c>.

**References**

Andrello, M., Darling, E. S., Wenger, A., Suárez‐Castro, A. F., Gelfand, S., & Ahmadia, G. N. (2022). A global map of human pressures on tropical coral reefs. Conservation Letters, 15(1), e12858.

Ayre, D. J., & Hughes, T. P. (2000). Genotypic diversity and gene flow in brooding and spawning corals along the Great Barrier Reef, Australia. Evolution, 54(5), 1590–1605.

Beyer, H. L., Kennedy, E. V., Beger, M., Chen, C. A., Cinner, J. E., Darling, E. S., ... & Hoegh‐Guldberg, O. (2018). Risk‐sensitive planning for conserving coral reefs under rapid climate change. Conservation Letters, 11(6), e12587.

Cowen, R. K., Paris, C. B., & Srinivasan, A. (2006). Scaling of connectivity in marine populations. Science, 311(5760), 522–527.

Gotelli, N. J. (1991). Metapopulation models: The rescue effect, the propagule rain, and the core-satellite hypothesis. The American Naturalist, 138(3), 768–776.

Greiner, A., Andrello, M., Darling, E., Krkošek, M., & Fortin, M. J. (2022a). Limited spatial rescue potential for coral reefs lost to future climate warming. Global Ecology and Biogeography, 31(11), 2245-2258.

Holstein, D. M., Paris, C. B., & Mumby, P. J. (2014). Consistency and in- consistency in multispecies population network dynamics of coral reef ecosystems. Marine Ecology Progress Series, 499, 1–18.

Kool, J. T., Paris, C. B., Barber, P. H., & Cowen, R. K. (2011). Connectivity and the development of population genetic structure in Indo-West Pacific coral reef communities. Global Ecology and Biogeography, 20(5), 695–706.

Manel, S., Loiseau, N., Andrello, M., Fietz, K., Goñi, R., Forcada, A., ... & Mouillot, D. (2019). Long-distance benefits of marine reserves: myth or reality?. Trends in Ecology & Evolution, 34(4), 342-354.

Mouquet, N., & Loreau, M. (2003). Community patterns in source-sink

metacommunities. The American Naturalist, 162(5), 544–557.

Pebesma E, Bivand R (2023). Spatial Data Science: With applications in R. Chapman and Hall/CRC. doi:10.1201/9780429459016, https://r-spatial.org/book/.

Pebesma E (2018). “Simple Features for R: Standardized Support for Spatial Vector Data.” The R Journal, 10(1), 439–446. doi:10.32614/RJ-2018-009, <https://doi.org/10.32614/RJ-2018-009>.

Saura et al., 2014

UNEP-WCMC and IUCN (2024), Protected Planet: The World Database on Protected Areas (WDPA) and World Database on Other Effective Area-based Conservation Measures (WD-OECM) [Online], April 2024, Cambridge, UK: UNEP-WCMC and IUCN. Available at: [www.protectedplanet.net](http://www.protectedplanet.net).

Vogt‐Vincent, N. S., Mitarai, S., & Johnson, H. L. (2023). High‐frequency variability dominates potential connectivity between remote coral reefs. Limnology and Oceanography, 68(12), 2733-2748.

Wood, S., Paris, C. B., Ridgwell, A., & Hendy, E. J. (2014). Modelling dispersal and connectivity of broadcast spawning corals at the global scale. Global Ecology and Biogeography, 23(1), 1-11.

**Supplementary Materials 2**. Networks of the Refugia

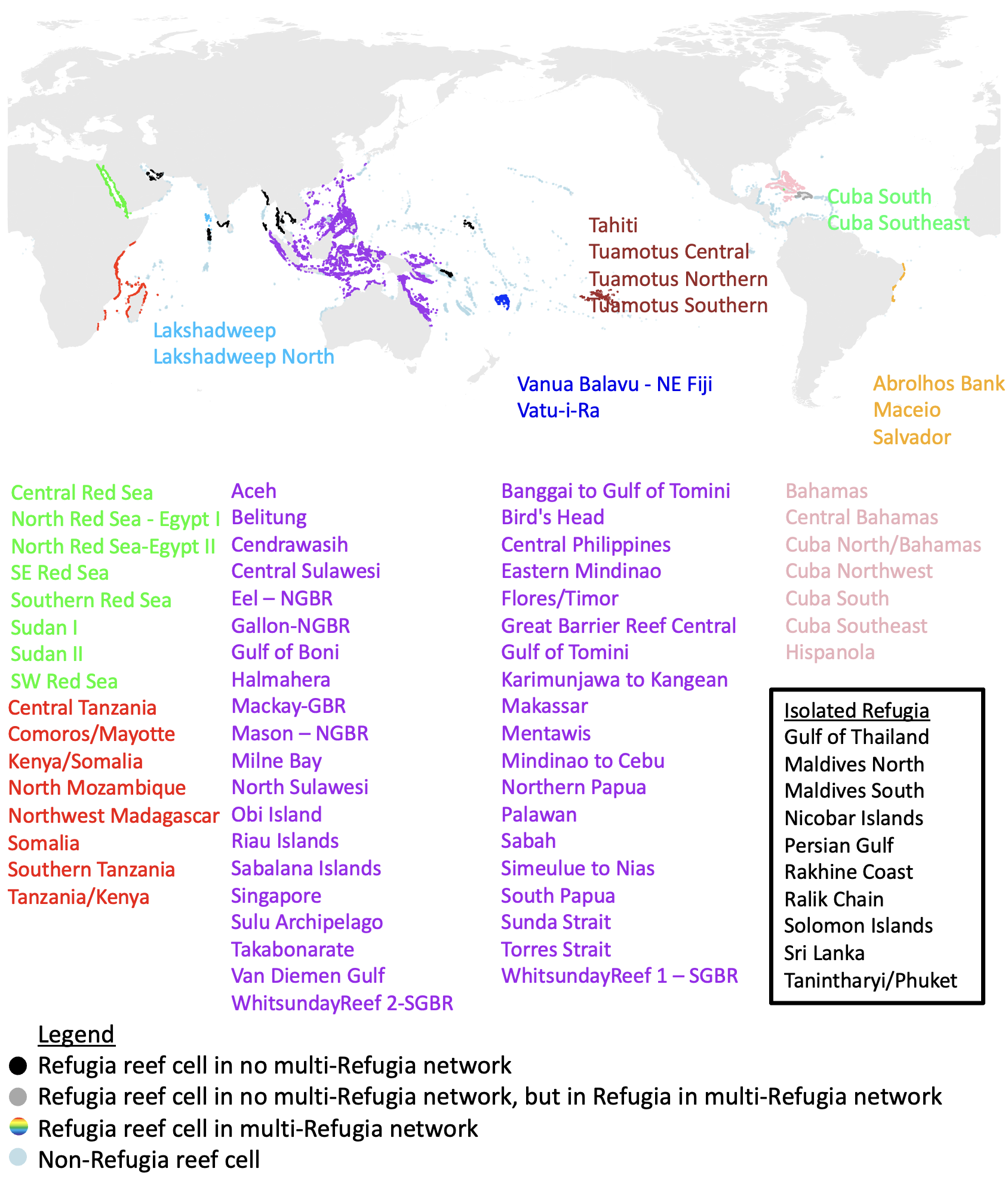

**Figure 1**. *Original Reef Networks of the 83 Refugia* - The map shows the connectedness of the 82 Refugia identified by Beyer et al. (2018) analysis. Refugia reef cells are coloured according to what multi-Refugia network at least some of those Refugia reef cells are contained within when all reefs are intact (reef cells in dark grey are the other reef cells in the Refugia in multi-Refugia networks); the colour of the reef cells on the map corresponds with the colour of the names of the Refugia. The Refugia listed in the box (black text) are isolated Refugia that are not connected to other Refugia through larval connectivity. Reef cells shown in light blue are not in Refugia, but some are in some of the same networks as the Refugia.

**Supplementary Materials 3**. Status of Refugia

The 83 Refugia are in 56 different reef networks when all reef cells are intact (Fig. 3a main text). Nine of those reef networks contain Refugia reef cells from more than one Refugia (multi-Refugia networks) and the other 47 only contain Refugia reef cells from one Refugia. Those nine multi-Refugia networks contain 73 Refugia (3,760 reef cells, 1,218,240 km^2^); the remaining ten Refugia are not in multi-Refugia networks (Fig. 3a main text).

When all non-refugia reef cells are lost, the number of reef networks containing Refugia increases from 56 to 123. Also, only 66 Refugia (3,388 reef cells, 1,097,712 km^2^) remain in multi-Refugia networks. Nineteen Refugia maintain all connections with other Refugia (Fig. 1a), 47 Refugia lose some of their connections with other Refugia (Partially Disconnected Refugia; Fig. 1b) and seven Refugia lose all connections with other Refugia (Disconnected Refugia; Fig. 1c) (Fig. 3b).

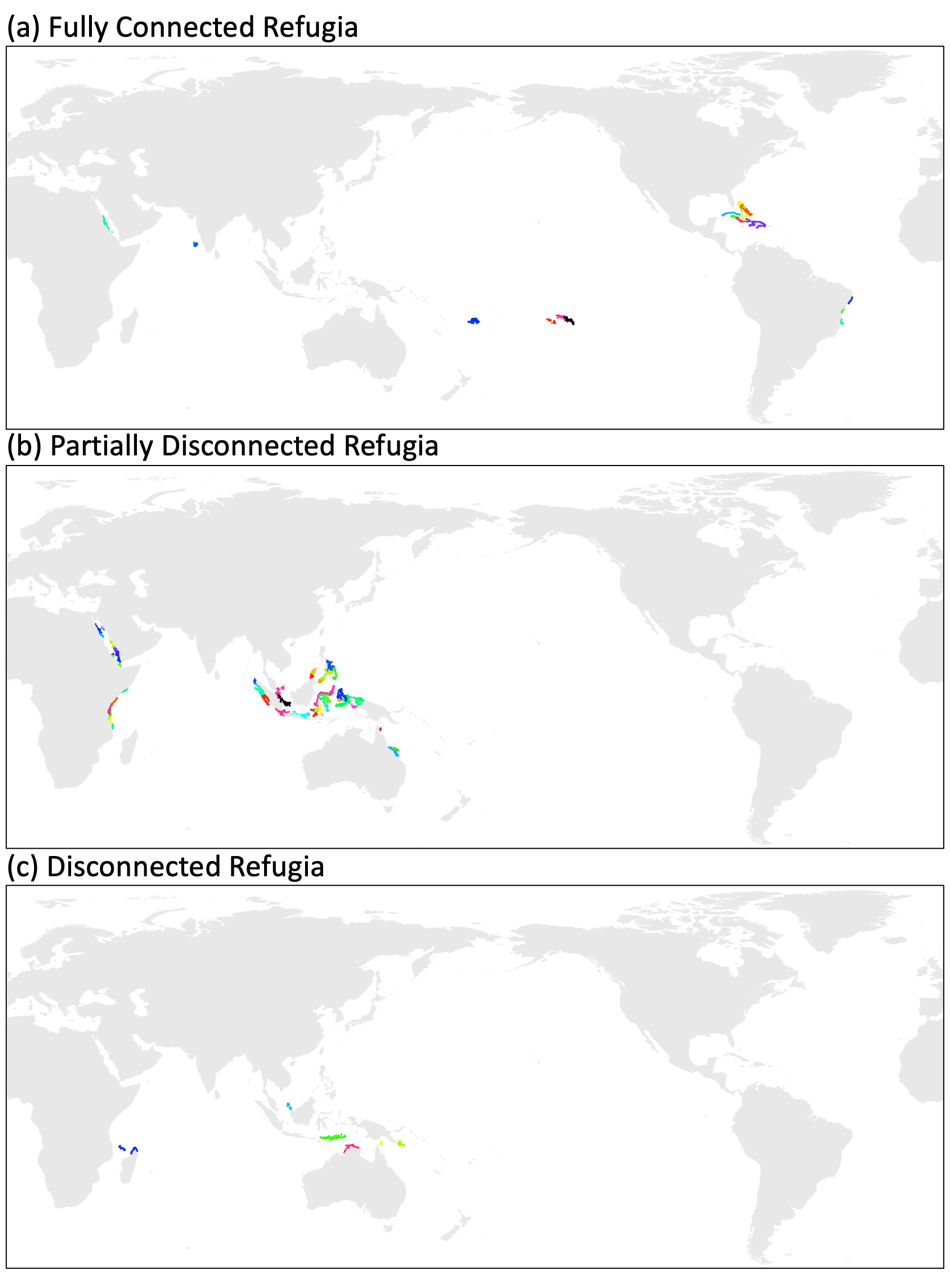

**Figure 1**. *Status of Refugia when Only Non-Refugia Reef Cells Remain* - This map shows Refugia reef cells coloured according to which reef network they are in, same colour means same network. Whether a Refugia reef cell is pictured in (a), (b), (c) is based on whether their Refugia maintains connections with other Refugia when only non-refugia reef cells remain.

When the 10 stepping stone reef cells are added back in (Fig. 2 main text), the reef networks regain some connections. Specifically, the 83 Refugia are now in only 113 different networks (Fig. 3c main text). Twenty seven refugia retain all present-day connections with other Refugia (Fig. 2a), 42 Refugia are now Partially Disconnected Refugia (Fig. 2b) and only 4 are Disconnected Refugia (Fig. 2c).

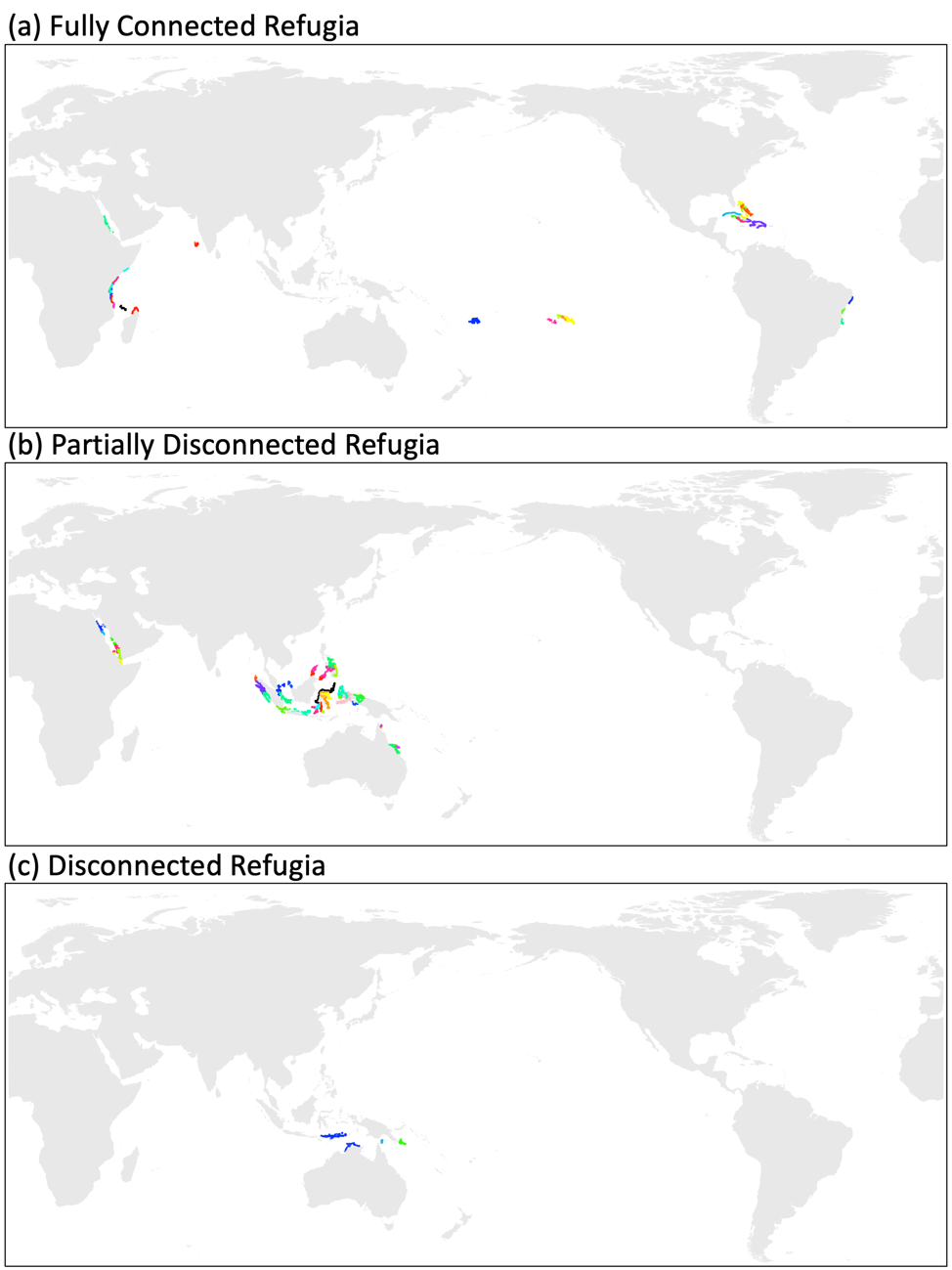

**Figure 2**. *Status of Refugia when Stepping Stones Also Maintained* - This map shows Refugia reef cells coloured according to which reef network they are in, same colour means same network. Whether a Refugia reef cell is pictured in (a), (b), (c) is based on whether their Refugia maintains connections with other Refugia when stepping stone non-refugia reef cells are the only non-refugia reef cells maintained.

When the 6 ‘connector’ stepping stone reef cells are added back in (Fig. 2 main text), the reef networks regain some connections. Specifically, the 83 Refugia are now in only 114 different networks (Fig. 3). Nineteen refugia retain all present-day connections with other Refugia (Fig. 4a), 50 Refugia are now Partially Lost Refugia (Fig. 4b) and only 4 are Lost Refugia (Fig. 4c).

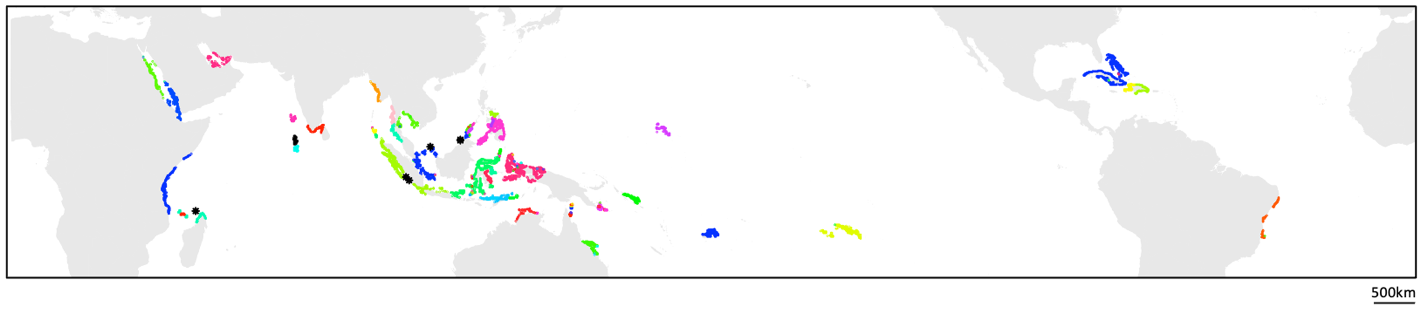

**Figure 3**. *Refugia reef cells + connector stepping stones, 83 Refugia in 114 networks* - Bright coloured reef cells indicate Refugia reef cells, coloured according to which reef network they are in (same reef network, same colour). Black stars indicate stepping-stone reef cells.

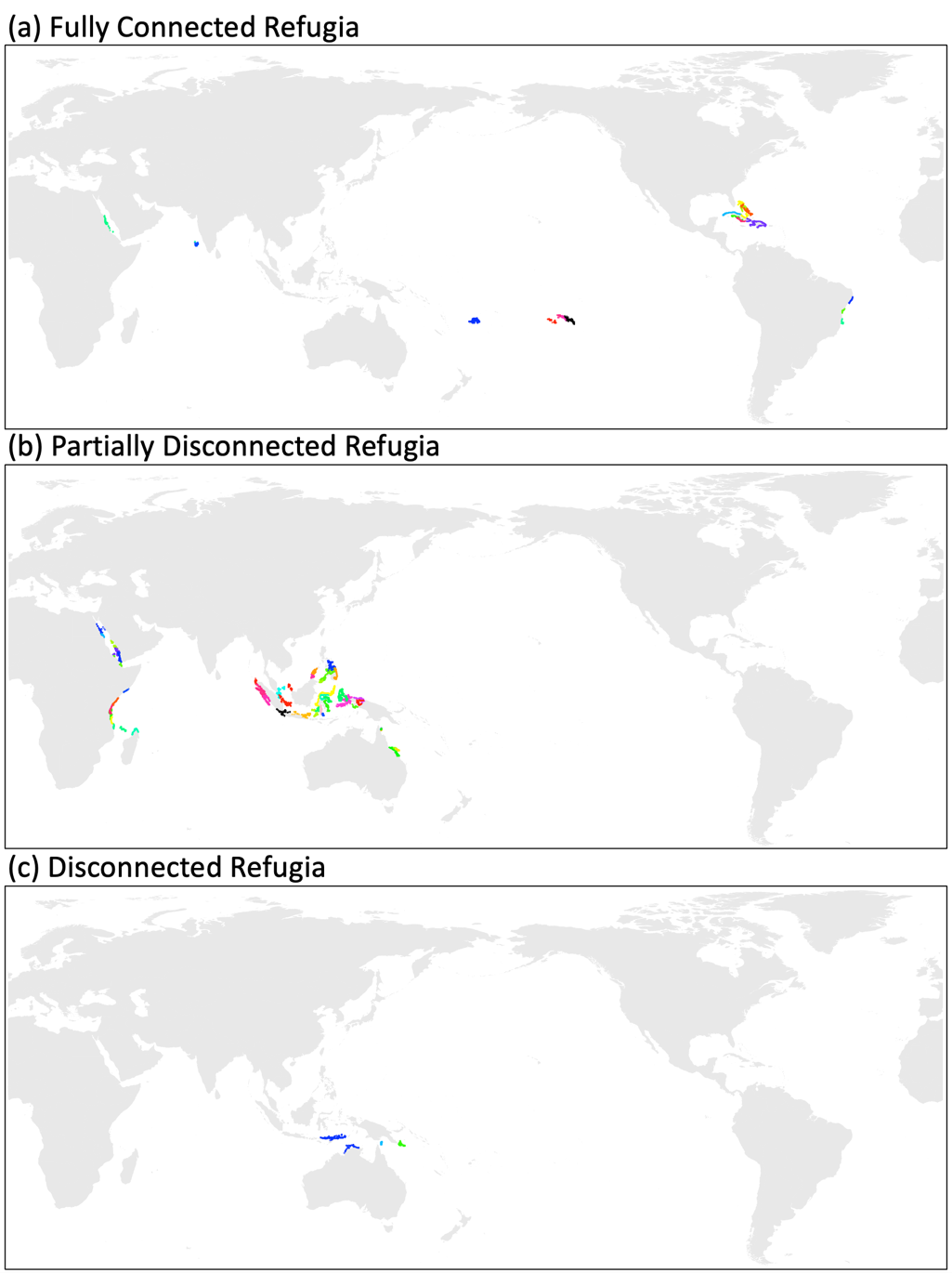

**Figure 4**. *Status of Refugia when Connected Stepping Stones Also Maintained* - This map shows Refugia reef cells coloured according to which reef network they are in, same colour means same network. Whether a Refugia reef cell is pictured in (a), (b), (c) is based on whether their Refugia maintains connections with other Refugia when stepping stone non-refugia reef cells are the only non-refugia reef cells maintained.

**Supplementary Materials 4**. Imperfect Overlap

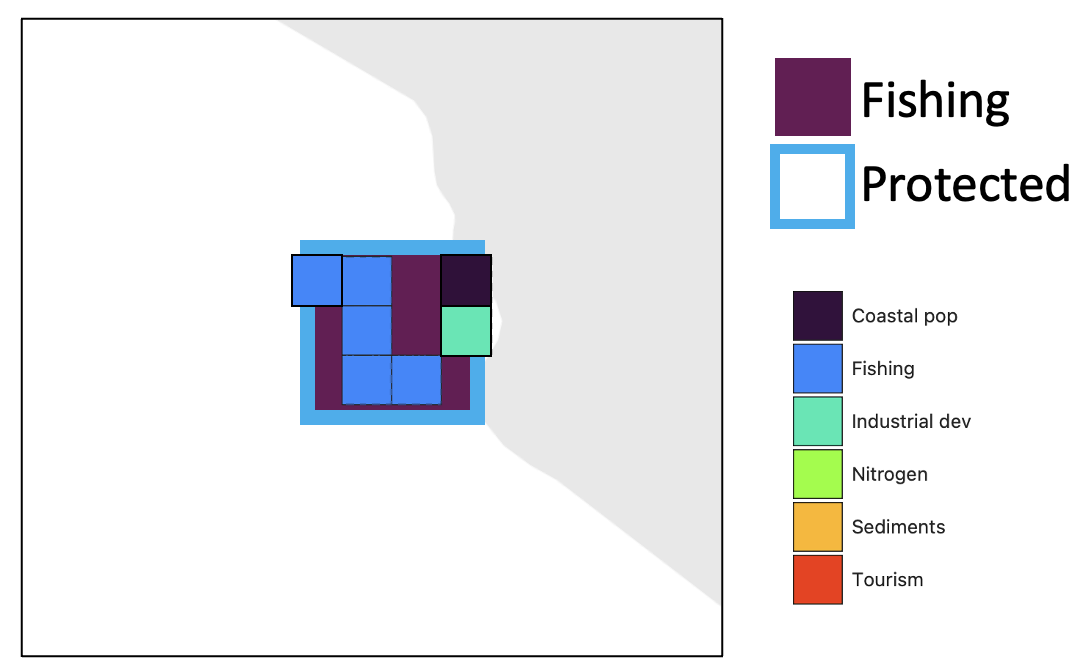

**Figure 1**. *Imperfect Overlap Between Grid Cell Types* – The larger purple grid cell is derived from Wood et al. (2014)’s grid cells (and corresponds with stepping stone Indonesia2; also see Figure 5 in the main text) and the smaller grid cells are derived from Andrello et al. (2022)’s grid cells. As described in Figure 5 in the main text, sometimes the stepping stone grid cells overlapped with multiple Andrello et al. (2022) grid cells. This was the one instance where all of the Andrello et al. (2022) grid cells that overlapped with a stepping stone grid cell did not have the same Top Anthropogenic Pressure. In this case, since most of the Andrello et al. (2022) grid cells that overlapped with Indonesia2 had ‘Fishing’ as their top pressure, we classified the top pressure of Indonesia2 as ‘Fishing’.

**Supplementary Materials 5**. Paired Stepping Stones

We repeated the same analysis described in the main text for paired sets of non-refugia reef cells to explore whether maintaining pairs of non-refugia reef cells as paired stepping stones would enable further re-connection of Refugia reef networks. When we did so, we found that no additional Refugia regained connections with other Refugia (i.e. the number of Fully Connected Refugia, Disconnected Refugia and Partially Disconnected Refugia was the same as when only the 10 stepping stones described in the main text were maintained) but that the number of reef networks containing Refugia decreased from 113 to 107 when the paired stepping stones were added (see Fig. 1 below).

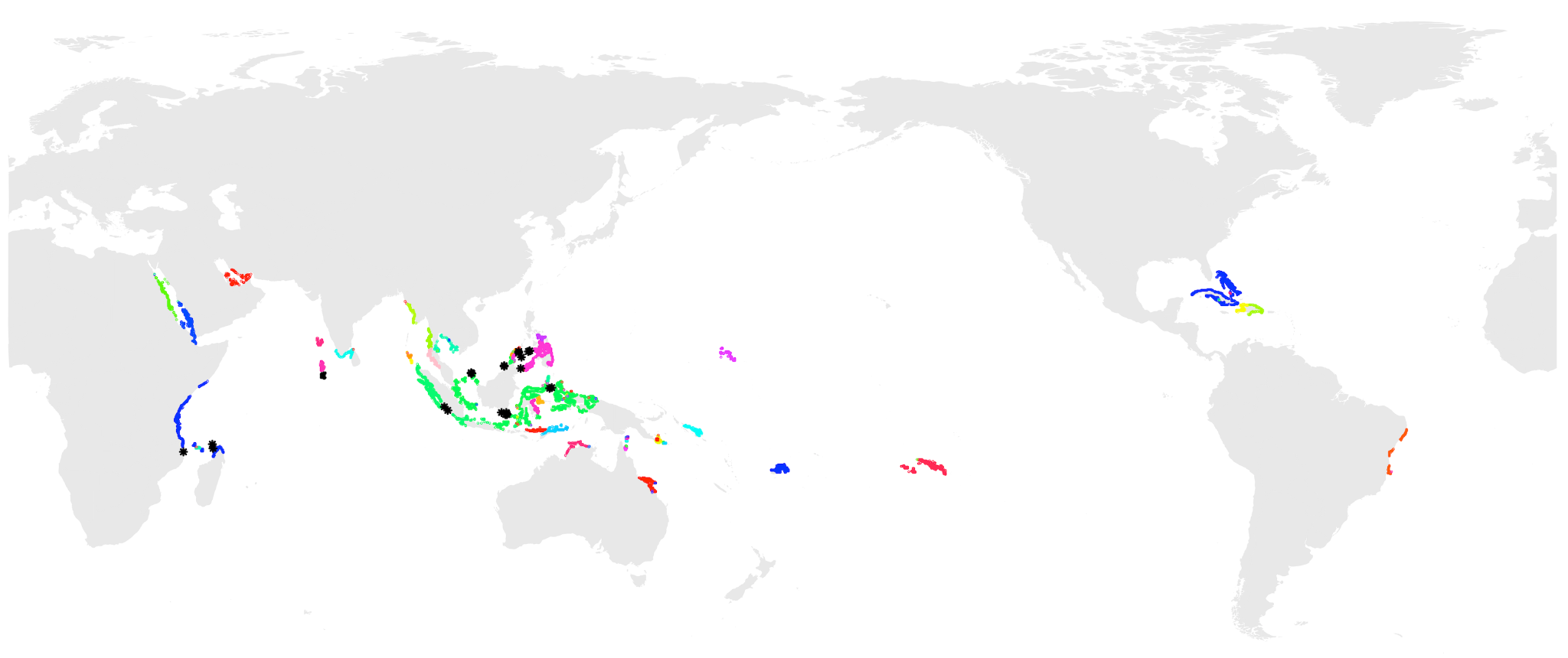

**Figure 1**. *Refugia Networks with Paired and Individual Stepping Stones* – This map shows Refugia reef cells coloured according to which reef network they are in when the individual stepping stones (discussed in the main text and shown in Figure 2) and paired stepping stones are maintained. Individual stepping stones and paired stepping stones shown as black grid cells in the map above. Reef cells shown in the same colour are in the same reef network.

When we start from the 6 connector stepping stone reef cells only and explore adding paired connector stepping stone reef cells, we find 4 paired connector stepping stone reef cell sets. Once again, no additional Refugia regained connections with other Refugia (i.e. the number of Fully Connected Refugia, Disconnected Refugia and Partially Disconnected Refugia was the same as when only the 10 stepping stones described in the main text were maintained). The number of reef networks containing Refugia decreased from 114 to 112 when the paired connector stepping stones were added (see Fig. 2 below).

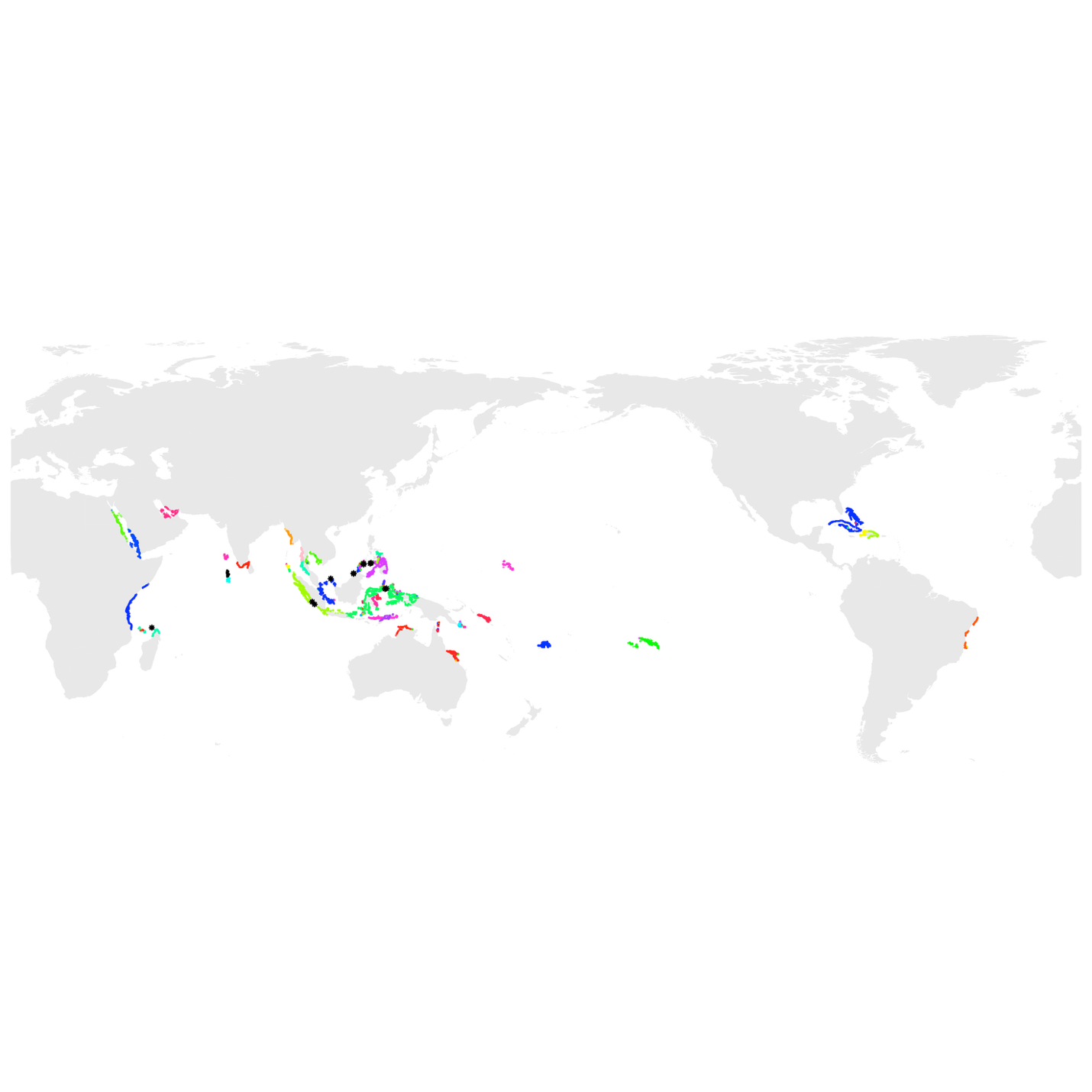

**Figure 2**. *Refugia Networks with Paired and Individual Connector Stepping Stones* – This map shows Refugia reef cells coloured according to which reef network they are in when the individual connector stepping stones (discussed in the main text and shown in Figure 2) and paired connector stepping stones are maintained. Individual connector stepping stones and paired connector stepping stones shown as black grid cells in the map above. Reef cells shown in the same colour are in the same reef network.
